# Multi-step femtosecond laser-fabricated membranes for regulated migration of biomolecules and cells

**DOI:** 10.64898/2026.05.22.726371

**Authors:** Anupam Gopalakrishnan, Akhila J. Denduluri, Shantae Gallegos, Izaiah Ramirez, Stephanie E. Schneider, Zeynep Çetinkaya, Heiko Kabutz, Alexander Hedrick, Kaushik Jayaram, Corey Neu, Gregory L. Whiting

## Abstract

Organ-on-chip (OoC) systems enable the recapitulation of key structural and functional characteristics of human tissues within controlled micro-engineered environments. In mechanically active tissues such as musculoskeletal, cardiac, and vascular systems, the incorporation of dynamic physical forces is essential for replicating the biomechanical cues governing cellular morphology and functional responses *in-vivo*. Without such stimuli, OoC models may fail to capture physiologically relevant tissue behaviors. Porous and semi-permeable membranes are critical components of OoCs, facilitating selective transport of nutrients, gases, and signaling molecules between cellular compartments to support biologically accurate barrier replication. Hence, fabrication strategies that permit precise modulation of membrane permeability are desirable to accommodate for the varying needs in pore size and porosity across organ systems. This study presents a two-stage fabrication process for stretchable, microporous polydimethylsiloxane (PDMS) membranes using femtosecond (fs-) pulse laser drilling. The laser-ablated pores exhibit a characteristic conical morphology, with diameters tapering from the laser entry to exit point. By modulating laser power and number of pulses, 6-15 μm exit-end pore diameters were achieved in 50 μm thick PDMS films. The membranes demonstrated strong mechanical resilience, with a 5–12% reduction in Young’s modulus after 500 cycles of strain loading. Furthermore, membranes fabricated at lower laser powers exhibited superior retention of elasticity, highlighting the influence of processing parameters on mechanical behavior. Cytocompatibility and permeability assessments confirmed that the membranes supported sustained cell viability and proliferation over at least three days. In size-restricted membrane pore geometries, cellular migration was constrained without any inhibition of biomolecular transport. This selective permeability is critical in multilayer OoC architectures, where a balance between biomolecular diffusion and cellular compartmentalization is necessary to preserve distinct tissue interfaces and functional organization. This work presents fs-laser micro-drilling as a robust and tunable fabrication strategy for producing mechanically resilient, selectively permeable PDMS membranes for physiologically relevant OoC applications.

## 1. Introduction

Organ-on-chip (OoC) systems are micro-engineered platforms that replicate key structural, functional, and physiological features of human organs *in vitro*, offering a compelling alternative to traditional cell culture and animal models for drug testing and disease research [1–3]. Typically, OoCs constitute of microfluidic channels, cells, extracellular matrix (ECM) components, and integrated sensors to recreate functional, mechanical and biochemical gradients to mimic organ-level responses [4,5]. To function effectively, these systems require cell-friendly surface chemistries, must be biocompatible, support fluidic manipulation, and enable precise control of the microenvironment [1,6]. Despite rapid progress, significant challenges remain, including the complexity of reproducing multi-tissue, inter-compartmental interactions, achieving long-term stability, standardizing fabrication techniques to ensure scalability and reproducibility, and function-optimized, minimally-obtrusive sensor integration into the platform [2,7,8].

One of the most functionally crucial elements of OoC systems are semipermeable membranes, where controlled permeability governs tissue function and homeostasis. These porous membranes enable the selective transport of nutrients, gases, and signaling molecules while maintaining compartmental separation between distinct cell populations. Custom semipermeable membranes can be engineered to replicate physiological barriers such as the blood–brain barrier, alveolar interface, and gut epithelium [3,9,10]. Additionally, the design and fabrication of membrane porosity, including pore size, density, and spatial distribution, remains a critical and technically challenging aspect of achieving realistic organ-level function in OoC systems [10,11]. The fabrication of thin porous membranes often involves expensive microfabrication infrastructure or track-etching methods, which typically sacrifice pore position control and optical transparency [11,12].

Many human organs and tissues are subject to continuous biomechanical stimulation *in vivo*, such as the rhythmic expansion and contraction of the lung during respiration, the pulsatile mechanical forces within the vasculature, and the compressive and shear forces experienced by articular cartilage and synovial membranes within knee joints [4,13–16]. Faithful reproduction of these forces within OoC systems is essential for eliciting physiologically relevant cell behavior and barrier function [4,9]. Accordingly, OoC platforms modelling such tissues commonly incorporate pneumatically actuated flexible membranes, deformable channel walls, or compression-loading mechanisms that impose cyclic mechanical strain on cultured cells, a capability that demands material architectures with well-defined, tunable, and stable elastomeric properties [7,13,17–19]. However, many OoCs have excluded mechanical stimulation due to the additional complexities introduced by device deformation. Materials used in mechanically active regions must be capable of elastic deformation at physiologically relevant strain levels while preserving structural integrity and bonding with the surrounding architecture over thousands of loading cycles to appropriately model *in vivo* conditions [20,21].

In addition, these materials must demonstrate resistance to fatigue and wear to ensure device integrity and consistent loading performance throughout the experimental timeline. Consequently, careful consideration must be given to both the mechanical properties and fabrication of the membranes. In applications where mechanical stimulation is critical, PDMS is frequently employed because its stiffness can be tuned to mimic target biological environments and its elasticity enables reversible deformation under applied loads [11,22]. Given this, novel fabrication techniques that can produce membranes with controlled pore geometry are increasingly being adopted in biomedical engineering. Commercial track-etched membranes remain widely used in static OoCs as they are low-cost, readily available, and offer well-defined pore diameters [23,24]. However, they are generally made from rigid polymers like polyethylene terephthalate or polycarbonate, making them incompatible with elastomeric devices designed to reversibly deform [12,25]. In stretchable PDMS films, manufacturing pores with sub-10 μm diameters at high densities is commonly achieved via soft lithography-based microfabrication in which micropillar features are defined by photolithography or deep reactive ion etching and then transferred through molds [26,27]. Although micro-molding offers high resolution and precise control over feature dimensions, the manual processing steps involved can introduce pattern distortion, membrane warping, and output variability that ultimately compromises reproducibility. Additionally, the reliance on clean rooms can make these methods difficult to implement in biology-focused labs. The fixed mold-based designs act as a constraining factor for rapid iteration of pore geometry and density when optimizing for specific biological applications [28,29]. Direct dry etching of PDMS templates has also been explored with moderate success, but the process is hindered by challenges such as surface roughness, scalability, and template thickness [30–32]. These limitations highlight the need for membrane fabrication approaches that offer mechanical compliance for dynamic loading, precise control over pore geometry, and independence from mold-based designs [11,22,33]. In this regard, ultrafast laser micromachining has emerged as a promising alternative for fabricating microfluidic devices and porous membranes [3,34,35].

Laser machining of dielectric materials, such as PDMS, follows a photothermal ablation mechanism initiated by multi-photon absorption and impact ionization. The energy absorbed during these processes generates an electron avalanche, resulting in melt-ejection of material once the ablation fluence threshold is exceeded [33,36]. Consequently, manipulating the thermal dynamics of the ablation zone enables precise tuning of pore geometries and surface properties such as wettability and reflectance [37–39]. Control over geometry through thermal regulation has previously been demonstrated through optimization of intrinsic laser parameters, such as spot size [40], wavelength [41], pulse width [42], pulse energy [35,43] and pulse repetition frequency (PRF) [44]. For instance, increasing pulse energy typically broadens feature dimensions [35,43], whereas decreasing the PRF tends to reduce them [44]. Such intrinsic parameters may require complex optical equipment to control and are fundamentally limited by the laser system’s hardware. Relying solely on intrinsic parameters also introduces trade-offs, such as dropping below the ablation threshold when attempting to minimize the heat-affected zone [45,46]. On the other hand, modulating operational parameters provides a more flexible alternative to regulate thermal dynamics and optimize fabrication without significantly altering the fundamental laser configuration. Despite this potential, investigations so far have been limited primarily to spatial and environmental factors such as laser scan speed and ambient temperature [44,47]. This leaves a significant gap in managing transient physical phenomena during fabrication, particularly, localized heat accumulation and plasma plume shielding, where ionized vapor absorbs incoming radiation [48–50].

To mitigate the thermal damage arising from laser-induced patterning, we introduce a two-stage ablation approach that incorporates a short temporal pause within the laser pulse sequence, allowing the plasma plume to dissipate between exposures. An ultraviolet (UV) femtosecond-pulsed laser (fs-laser) (λ = 343 nm, pulse width = 247 fs) was used to drill micron-sized holes for the fabrication of porous PDMS membranes. The laser pulse train was delivered in two stages, separated by a brief (10 s) delay, which further reduced the thermal damage to the area surrounding the ablation site. The resulting membranes preserve their intended mechanical properties, remain cytocompatible, and effectively minimize cell migration while permitting biomolecular transport. This method offers a reliable technique for producing tunable microporous membranes suitable for integration into dynamic OoC systems.

## 2. Materials and Methods

### 2.1. PDMS membrane fabrication

Thin films for the membranes were obtained by spin-casting PDMS (Sylgard 184) onto 75x25 mm glass microscope slides pre-treated with Trichloro(3,3,3-trifluoropropyl) silane. Glass slides were cleaned in a plasma chamber for 5 mins and placed in a desiccator under vacuum along with silane for 1 hr (40 µL of silane per glass slide). PDMS was prepared by thoroughly hand-mixing the base and curing agent in 10:1 weight ratio. To remove the air bubbles formed while stirring, the mixture was placed in a desiccator under vacuum for 20 minutes. The mixture was then spin-cast onto the silanized glass slides at 1200 rpm for 60 sec and immediately cured at 100°C for 40 min. This procedure resulted in PDMS films that were 50 µm thick. Subsequently, the PDMS films, while on their glass substrates, were used for laser micromachining.

### 2.2. Laser microfabrication

An ultraviolet (UV) fs-laser (CARBIDE-CB5, Light Conversion, Lithuania) with a pulse duration of 247 fs, wavelength of 343 nm and a spot size of 8-10 μm was used to fabricate the PDMS membranes. The pulse repetition frequency (PRF) of the laser can be varied from 100 kHz to 300 kHz. The corresponding output pulse energy and power generated, interchangeable with the selected PRF, are shown in Table 1. Control of the laser system, including selection of PRF and number of pulses per pore (NP) was achieved using DMC software (Direct Machining Control, Lithuania).

**Table 1.**
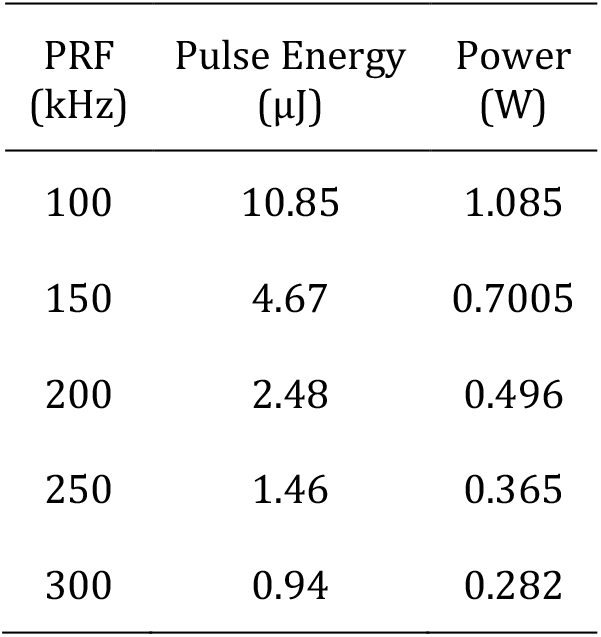
The output laser pulse energy and power at various pulse repetition frequencies (PRFs).

Membranes were fabricated using two laser drilling protocols: single-stage and two-stage. In the single-stage protocol, pores were drilled using three independent pulse trains at PRFs of 150, 200 and 250 kHz, each consisting of 40 pulses (NP = 40). In the two-stage protocol, the 40-pulse train was split equally into two sequential stages of 20 pulses each, separated by a brief delay (∼10-15 s) accounting for the time required for the laser system to switch between PRFs states. The scan speed for both protocols was set at 100 mm/s. All membrane outlines were cut in a single stage by repeatedly scanning each line 35 times at a PRF of 250 kHz and a scan speed of 500 mm/s. After fabrication, the PDMS membranes were ultrasonically washed in an ethanol bath and then peeled off the glass slide for subsequent characterization. The entire laser microfabrication process was carried out in air at ambient temperature (20 ± 2 °C) and humidity (30 ± 10%). The CAD models for the membrane perimeters were designed in SolidWorks (Dassault Systèmes) and imported to DMC software, where a point-based hatching pattern was generated within these boundaries to define the specific coordinates for laser drilling. A uniform center-to-center pore pitch of 40 µm was selected for all fabricated samples used in characterization. Each data point was obtained by averaging values from 3 replicate samples.

### 2.3. Imaging

Membrane pores were imaged under a laser confocal microscope (VK-X3000 Surface Profiler, Keyence Corporation of America). The root mean square surface roughness (S_q_) of the entry surface was measured from the obtained data using Multifile Analyzer Software (Keyence Corporation of America). ImageJ software was then used to measure the roundness, circularity, and area of pores on both the entry and exit surfaces. Effective entry and exit pore diameters were calculated from the measured pore areas by assuming perfectly circular pore ends.

### 2.4. Tensile testing

Mechanical characterization of the fabricated membranes was performed via uniaxial tensile testing using an MTS Insight 2 mechanical testing machine equipped with a 100 N load cell. Rectangular film samples with a length of 100 mm, gauge length of 50 mm and a width of 5 mm were used. The samples were subjected to a loading of 500 cycles of sinusoidal strain at an amplitude of 30% at 0.5 Hz, where required. Tensile failure tests were performed before and after cycling at a constant strain rate of 1 mm/s. The stress-strain curves were recorded and averaged across three independent replicates. The value of elastic modulus was calculated from the slope of the stress-strain curve within the linear regime (<10% strain).

### 2.5. Statistical Analysis

We evaluated three characteristics of the fabricated membranes: exit diameter, entry diameter and entry surface roughness, with the latter serving as a metric for thermal damage. Through preliminary trials, the pulse repetition frequency (PRF) of the laser and the number of incident pulses (NP) were identified as the main factors governing these features. To further investigate the effects of these parameters in a two-stage process, PRF and NP for each of the two stages in the pulse train (I-PRF, I-NP, II-PRF and II-NP) were set to 4 levels to implement a 4-factor-4-level experimental design. Table 2 shows the values and levels for the processing parameters and Table S1 shows the parameter combinations evaluated. JMP^®^ (SAS Institute Inc., Cary, NC, USA) software was used to generate the experimental design, conduct analysis of variance (ANOVA) shown in Table S2, and develop the predictive model.

**Table 2.**
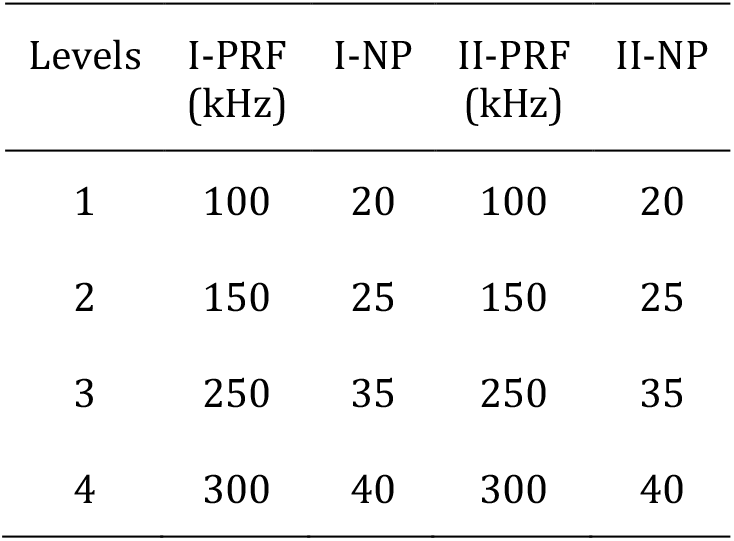
Laser parameters and their levels used for experimental design.

### 2.6. Atomic Force Microscopy

The PDMS membrane surfaces were scanned and indented using an atomic force microscope (DriveAFM, Nanosurf AG, Switzerland) equipped with a silicon pyramidal probe tip (HQ:NSC15/Al BS, Bulgaria). The Young’s moduli at various locations around the pores were determined by fitting the force-distance curves obtained from nanoindentation. The fitting was performed on the software provided by the AFM manufacturer (Nanosurf Studio) using the Harding-Sneddon Cone model, the parameters for which are listed in Table 3. Young’s modulus measurements were taken at 3 or 5 locations for each sample. At each location, indentations were made in a 5×20 grid (pitch = 0.5 µm) lengthwise radially away from the pore edge, resulting in 5 sets of force-distance curves. Statistical comparisons between Young’s modulus at the pore edge (∼0.5 µm) and at ∼10 µm away from pore edge were performed using Welch’s independent samples t-test, which was selected to account for the unequal variances observed between the two measurements. Analyses were conducted separately for each sample using JMP statistical software. The significance threshold was set at α = .05. Effect sizes were calculated as Cohen’s d using the pooled standard deviation.

**Table 3.** Input values for the Harding-Sneddon Cone model used to fit force-distance curves obtained from nanoindentation.

| Parameter | Value |
| --- | --- |
| PDMS Poisson's ratio | 0.5 |
| Probe cone half-angle | 20° |
| Probe radius | 7 nm |
| Probe Young's modulus | 158 GPa |
| Probe Poisson's ratio | 0.28 |

### 2.7. Cell Viability

Membranes were submerged in 70% ethanol and washed via ultrasonication for 15 min to remove debris before transfer to non-treated TC plates (Falcon, 08-772-50). Transferred membranes were submerged in 70% ethanol and rinsed 3 times with Phosphate buffered saline (PBS) for sterilization before incubation with 20 µg/ml fibronectin (Sigma, F1141) in PBS at 37°C overnight and seeded with NIH 3T3s at 3,000 cells/cm^2^. NIH 3T3s were grown and maintained in Dulbecco’s Modified Eagle’s Medium (DMEM; High glucose with L-Glutamine; ATCC 30-2002), supplemented with 10% (*v*/*v*) fetal bovine serum (Gibco, A5670701), and 1% penicillin/streptomycin (1000 U/mL penicillin and 1000 µg/mL streptomycin; Gibco, 15140-122) at 37 °C and 5% CO_2_. Cells were passaged into a new 25 cm^2^ culture flask (Thermo Scientific, 156367) upon reaching 80% confluency. Viability was assessed via live/dead staining and confocal microscopy (Nikon A1R). 1 μl/ml calcein-AM (cal AM) (Invitrogen, C3100MP) and 2 μl/ml ethidium homodimer-1 (EthD-1) (Invitrogen, E1169) in PBS were added to samples and incubated at 37°C for 20 min prior to imaging. A Nikon 10× air objective (512×512 / acquisition matrix) and excitation wavelengths of 543 nm (Texas Red) and 488 nm (Fluorescein) were used for imaging. Images collected were processed in ImageJ (National Institutes of Health, Bethesda, MD), and viability was assessed using the percentage of live/dead cells.

### 2.8. Cell Transmigration

Fabricated PDMS membranes were bonded to the plastic components of membrane-free transwell inserts with the membrane’s exit side facing up towards the inside of the insert, where cells were seeded, following a standard transwell-style migration assay format [51]. To ensure a leak-free seal, membranes were bonded using a thin layer of uncured PDMS (Sylgard 184, 10:1 base:curing agent) around the edges avoiding the laser ablated pores, followed by curing at 80°C for 30 min. Inserts were sterilized in 70% ethanol, rinsed three times with sterile PBS, and coated with fibronectin (10 µg/mL, 100 µL per insert; Sigma-Aldrich) at 37°C for 24 hr [52]. NIH 3T3s (passage ≤ 50) were labelled with CellTrace™ CFSE (5 µM, 20 min at 37°C; Thermo Fisher) to identify cells on the membrane. Labelled cells were seeded into the upper chambers at 5,000 cells per insert (15,000 cells/cm^2^).

Two migration assays were conducted using membranes with 8 μm and 16 μm pore exit diameters. For each assay, the lower chambers were filled with 750 µL of one of three media conditions (n = 3 fabricated inserts per condition): serum-free A10 (negative control), A10 (control) or A10 with 10 ng/mL recombinant mouse PDGF-BB (PeproTech, 315-18) [53,54]. For each media condition, an unmodified FluoroBlok − insert (Corning, Cat. 351152) was included as a commercially available control. Plates were incubated for 24 hrs at 37 °C and 5% CO_2_ to allow cell transmigration through the membranes and then fixed in 4% PFA. Fixed CFSE-labelled cells were imaged by confocal microscopy using a 10× objective. For each well, five fields of view (top left, top right, bottom left, bottom right, and center) were acquired on (i) the seeding surface at the top of the membrane (non-migrated cells) and (ii) the surface where migrated cells accumulated. CFSE-positive cells were quantified using ImageJ (NIH). Images were thresholded using a consistent visual criterion to distinguish cells from background fluorescence, with thresholds applied uniformly across comparable conditions and adjusted only where necessary to account for differences in substrate background.

Images were despeckled to reduce noise prior to analysis. Particle analysis was performed using a size range of 15–5000 µm^2^ and circularity of 0.05–1.00 to exclude debris while retaining elongated NIH 3T3 fibroblasts. All analysis parameters were validated by visual inspection of representative images.

For each insert, the percentage of transmigrated cells, ***T***%, was calculated as

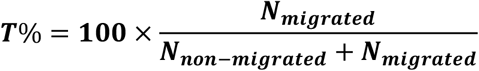

where ***N***_***migrated***_ **and *N***_***non***−***migrated***_ are the total numbers of CFSE-positive cells on the migrated and non-migrated surfaces, respectively. All conditions were tested in triplicate inserts per experiment unless otherwise noted.

### 2.9. Molecular Diffusion

Membranes were submerged in 70% ethanol and washed via ultrasonication for 15 min to remove debris before being placed between two channels in a microfluidic device during fabrication. Top and bottom channels were filled with 70% ethanol to reduce surface tension and rinsed with PBS. 2.5 mg/ml of 10 kDa (Sigma, FD10S) or 250 kDa (Sigma, FD250S) FITC-Dextran was dispensed into the top fluid channel with a 1 ml single-use syringe. Image acquisitions (6 frames per minute) of both 10 kDa and 250 kDa dextran in triplicate were acquired using a confocal microscope (Nikon A1R) in the 488 channel (for FITC), focused within the bottom fluid channel. The fluorescence Intensity was measured in ImageJ (NIH) and normalized against the maximum intensity value of the picture over 120 sec. Mean FITC fluorescence intensity was then obtained by averaging the values computed for n=3 devices and plotted as normalized mean intensity over time. To further characterize the difference in rate of diffusion between the two molecule sizes, the data was fit to a general exponential approach to equilibrium model using custom MATLAB code. An exponential model, defined as:

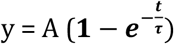

where A is the equilibrium value, and τ is the characteristic diffusion time, was used to fit the diffusion data to a general equation in MATLAB.

## 3. Results and Discussion

### 3.1. Membrane fabrication

The workflow for fs-laser fabrication of microporous PDMS membranes is illustrated in Figure 1A, alongside a representative image of a fabricated membrane (Figure 1B). In laser percussion drilling of PDMS, plasma-mediated ablation accompanied by melt ejection produces pores with a characteristic conical geometry. As illustrated in Figure 1C, this geometry manifests as a systematically larger diameter at the laser-incident (pore entry) surface relative to the opposing (pore exit) surface. This asymmetric geometry can be confirmed in the cross-sectional morphology (Figure 1D), with a gradual taper from entry to exit, reflecting directionally dependent energy deposition and material removal during the drilling process.

**Figure 1.**
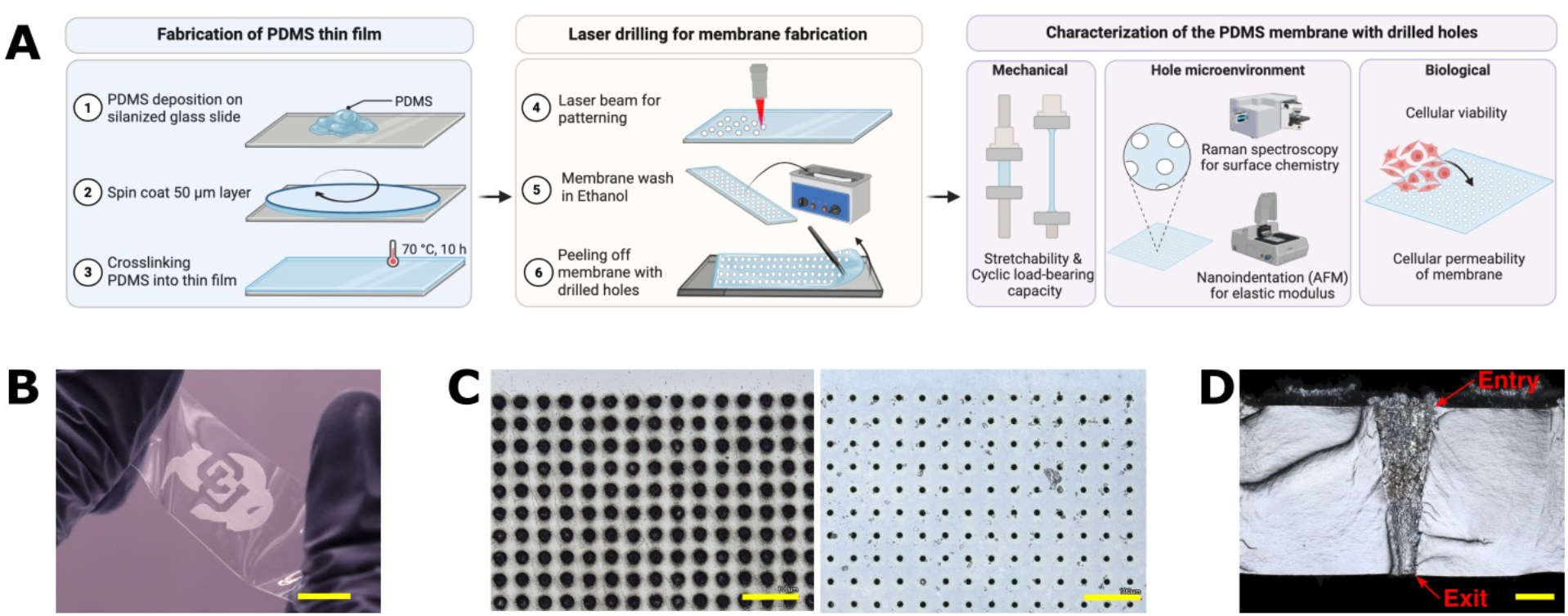
Fabrication workflow and morphological characterization of laser-drilled microporous PDMS membranes. (A) Schematic illustration of the PDMS membrane fabrication process – image created in Biorender. (B) Optical image of a 50 μm-thick PDMS membrane with the porous region (opaque) patterned in the shape of the University of Colorado Boulder logo (scale bar: 1 cm). (C-D) Confocal microscopy images of (C) the entry (left) and exit (right) surface of the porous membrane (scale bar: 100 μm). (D) A cross-section of a single pore showing a tapered geometry with a larger entry diameter and smaller exit diameter. (scale bar: 10 μm).

We investigated the effect of two fabrication parameters *i*.*e*., Pulse repetition frequency (PRF) and number of incident pulses (NP) on three key response variables: entry diameter, exit diameter, and entry surface roughness, with the latter serving as an indicator of thermally induced damage. For PRFs of 150, 200 or 250 kHz, the surface morphologies of PDMS membranes were fabricated using a single-stage pulse train (40 pulses) or a two-stage pulse train (2x 20 pulses) (Figure S1). In single-stage ablation, the variations in PRF exhibited an inverse relationship with pore size, whereby increasing PRF led to a reduction in both entry and exit diameters (Figure S2A) [35,44]. This trend is consistent with thermal effects and plasma shielding at higher repetition rates, which effectively limit material removal per pulse [41,55]. However, entry surface roughness exhibited only a weak dependence on PRF under single-stage ablation, with marginal reduction in surface roughness with increasing PRF. This could reflect the competing effects of thermally driven surface relaxation, which promotes smoothing [56–58] and plasma shielding, which introduces spatially non-uniform energy deposition and surface irregularities [59].

Switching the fabrication process from single-stage to two-stage ablation retained the previously observed inverse relationship between PRF and pore size. However, restructuring the pulse delivery into a two-stage sequence exhibited a stronger effect on surface roughness, whereby the surface roughness across all tested PRFs (Figure S2B) decreased, indicating a substantial mitigation of thermal damage. The implementation of the two-stage ablation reduced the entry surface roughness relative to the corresponding single-stage conditions. The difference in the surface roughness (reduction) between the two-stage and one-stage ablation conditions was 0.213, 0.210, and 0.177 μm at PRFs of 150, 200, and 250 kHz, respectively. Crucially, this improvement in surface quality was achieved without compromising pore dimensions relative to the corresponding single-stage ablation conditions (Figure S2A).

We corroborated these findings with microscopic observations, in which we noticed a marked reduction in crack formation at the entry surface for the two-stage process. This behavior may be related to the introduction of an inter-stage delay, which likely facilitates partial thermal relaxation and plume dissipation between the pulse trains [47–50]. The reduced persistence of the plasma plume likely minimizes shielding effects, enabling more efficient energy coupling in subsequent pulses while preventing excessive heat accumulation. These results demonstrate that temporal structuring of pulse delivery is an effective strategy for decoupling ablation efficiency from thermal damage in fs-laser drilling of PDMS membranes.

A four-factor, four-level experimental design and ANOVA analysis (observed results in Table S1, statistical analysis in Table S2) were employed to quantify the individual contributions of I–PRF, II– PRF, I–NP, and II–NP to variations in pore geometry and surface roughness, and to assess their statistical significance. The analysis confirmed that I–PRF and II–PRF were the dominant and statistically significant factors (p < 0.05) influencing the key response variables, indicating that PRF likely governs both pore dimensions and surface characteristics. The shared influence of these cofactors across multiple responses further provides a basis for examining interdependencies between outputs, such as the observed correlation between entry diameter and surface roughness. Entry diameter was strongly influenced by I–PRF and II–PRF, accounting for 45.25% and 41.24% of the total variance, respectively. A comparable trend was observed for entry surface roughness, where PRF again dominated (I–PRF: 51.53%; II–PRF: 35.49%), indicating that energy delivery rate is the primary determinant of both material removal and thermally induced surface modification at the laser beam-membrane interface. A positive correlation between entry diameter and surface roughness was observed, with larger entry diameters associated with increased roughness. This relationship suggests that conditions promoting enhanced ablation could also intensify local thermal effects, resulting in a more pronounced heat-affected zone. Such coupling is particularly relevant in the context of membranes fabricated for bio-interfacing, where surface topography can directly influence cellular behavior and attachment, biomolecule adsorption, and interfacial transport [60]. In contrast to entry diameter which seems to be mostly PRF-governed, the exit diameter was primarily governed by first-stage parameters, with I–PRF and I–NP contributing 41.03% and 30.11% of the variance, respectively. The second-stage parameters exhibited comparatively weaker, yet still meaningful, contributions. This distinction indicates that initial laser–material interactions largely determine pore breakthrough and through-material thickness of the pore geometry, while the second stage plays a more limited role in modifying exit dimensions. The stage-dependent influence of processing parameters provides a mechanism for independently tuning entry and exit pore features.

To demonstrate the tunability and versatility of the two-stage process, predictive modelling was used to identify parameter combination sets capable of achieving a constrained pore exit diameter (7–9 μm) while varying entry diameter (Table 4). Parameter combination sets were obtained to maximize, minimize, or normalize entry dimensions without deviating from the target exit diameter range. This capacity to independently tune entry and exit pore characteristics highlights the versatility of the two-stage fs-laser ablation process for designing porous membranes with application-specific transport and interfacial properties. The selected conditions were subsequently evaluated through mechanical testing, physicochemical surface characterization, and cell and biomolecular transport assays, thus enabling direct linkage between fabrication parameters, pore architecture, cellular compatibility and functional performance.

**Table 4.** Parameter combinations selected for characterization. Samples are labelled as Ø[Exit Diameter] - E_[Total Energy]_ with rounded values.

| Label | Step I |  | Step II |  | Total Pulse Energy Dosage (µJ) | Predicted Dia |  | Measured Dia |  |
| --- | --- | --- | --- | --- | --- | --- | --- | --- | --- |
|  | PRF (kHz) | NP | PRF (kHz) | NP |  | Exit (µm) | Entry (µm) | Exit (µm) | Entry (µm) |
| Ø8-E<sub>290</sub> | 100 | 20 | 150 | 20 | 290.93 | 8.65 | 29.57 | 8.18 | 28.39 |
| Ø8-E<sub>120</sub> | 150 | 20 | 250 | 30 | 121.25 | 7.67 | 23.89 | 7.99 | 20.75 |
| Ø8-E<sub>80</sub> | 200 | 25 | 250 | 25 | 81.10 | 7.81 | 20.93 | 7.90 | 19.68 |
| Ø16-E<sub>220</sub> | 150 | 40 | 250 | 40 | 218.82 | 16.71 | 25.46 | 16.14 | 24.71 |

PRF: Pulse Repetition Frequency, NP: Number of Pulses.

### 3.2. Tensile Characteristics

Mechanical robustness under physiologically relevant loading or deformation is a critical requirement for OoC platforms that aim to recapitulate movement-induced biomechanical cues, such as those present in lung alveolar or joint microenvironments [17,26]. To evaluate the mechanical behavior of the fs-laser fabricated PDMS membranes, uniaxial tensile testing was performed before and after cyclic loading. Membranes were subjected to 500 cycles of sinusoidal strain at 30% amplitude at 0.5 Hz, representing the upper range of physiological and pathological deformation reported for mechanically active tissues [61,62].

Representative optical images (Figure 2A) and corresponding exit-surface views (Figure 2B) demonstrate membrane deformation at 0% and 30% applied strain. No visible defects or discontinuities were observed under deformation, indicating that the membranes maintain structural integrity within the tested strain range. Across all the fabricated membranes tested, stress–strain responses (test set-up in Figure 2C) before-cyclic loading (Figure 2D) and after-cyclic loading (Figure 2E) exhibited an initial linear elastic regime up to approximately 10% strain, followed by a nonlinear regime at higher strains, consistent with elastomeric deformation behavior previously reported [63,64].

**Figure 2.**
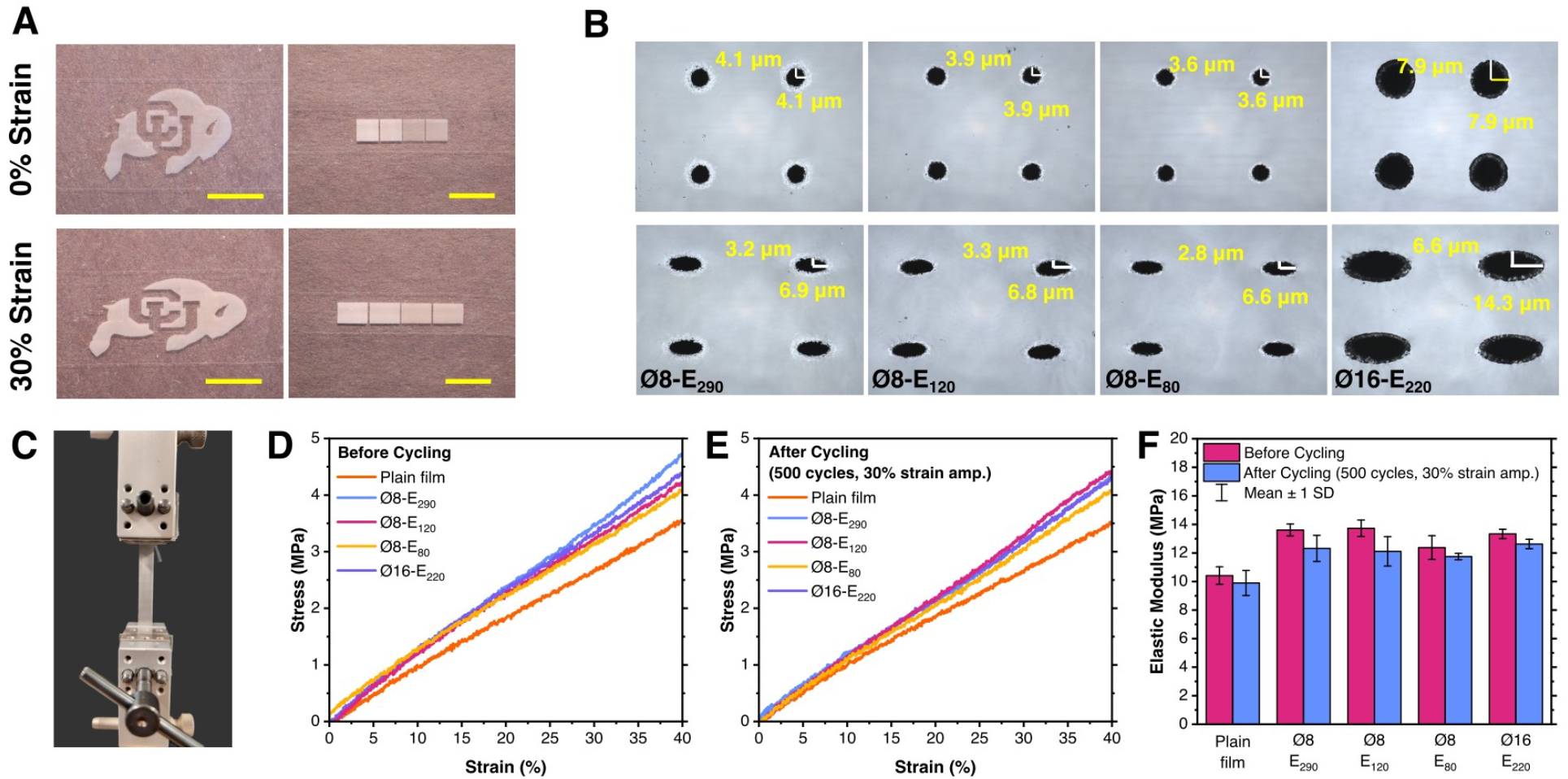
Laser-patterned PDMS membranes maintain structural integrity and exhibit stable mechanical properties under cyclic strain. (A) Optical images of patterned membranes under 0% (top) and 30% (bottom) strain (scale bar: 1 cm). (B) Confocal microscopy images of the exit surface of the microporous PDMS membranes fabricated with different laser parameters (Ø8-E_290_, Ø8-E_120_, Ø8-E_80_, Ø16-E_220_), under 0% (top) and 30% (bottom) strain. (C) The tensile test setup. The rectangular test specimen had a total length of 100 mm, a gauge length of 50 mm and a width of 5 mm. (D,E) Stress-strain behaviours of PDMS membranes (D) before and (E) after 30% cyclic tensile strain for 500 cycles at 0.5 Hz. (F) Elastic modulus of the PDMS membranes before and after cycling (n = 3).

Plain, non-patterned PDMS membranes exhibited an average modulus of 10.42 MPa, which is higher than the nominal bulk PDMS range (1–3 MPa). This variation could be due to thin-film effects associated with the spin-coating fabrication process. Specifically, elevated shear stresses during high-speed spin coating promote straightening of coiled polymer chains and increase effective crosslink density, resulting in enhanced stiffness relative to bulk material properties [65]. When compared to non-patterned films, PDMS membranes with fs-laser-drilled pores exhibited higher elastic moduli with values of 13.61 MPa (Ø8-E_290_), 13.73 MPa (Ø8-E_120_), 12.37 MPa (Ø8-E_80_), and 13.33 MPa (Ø16-E_220_) (Figure 2F). This increase in stiffness relative to plain PDMS could likely be due to thermally induced post-curing during laser ablation. As a thermosetting polymer, PDMS is susceptible to additional crosslinking when exposed to localized heating, resulting in a denser polymer network and enhanced mechanical stiffness. Despite variations in laser drilling parameters, the measured moduli remained within a narrow range, indicating minimal sensitivity of bulk mechanical properties of the membranes to the specific laser patterning conditions. This limited variation is likely due to the inherently low thermal conductivity of PDMS (∼0.27 W·m^−1^·K^−1^), which restricts heat transfer away from ablation sites and confines any thermal effects to highly localized regions. Consequently, the overall structural integrity and bulk mechanical response of the membrane remain largely unaffected. Minor differences in modulus are therefore attributed to secondary factors, including variations in membrane porosity (Figure S3) and differences in laser fluence. These findings suggest that, within the parameter space investigated, laser power and pulse number do not significantly influence the elastic modulus of the resulting membranes, consistent with previously reported observations in laser-processed PDMS systems [66].

Following cyclic tensile loading, all samples exhibited a reduction in elastic modulus in the range of approximately 5-12%. The post-cycling modulus values were measured at 12.32 MPa (Ø8-E_290_), 12.11 MPa (Ø8-E_120_), 11.74 MPa (Ø8-E_80_) and 12.62 MPa (Ø16-E_220_). In comparison, plain, non-patterned PDMS films exhibited a reduction of ∼5% under the same loading conditions. While previous studies have reported modest modulus reductions (∼2%) after fewer cycles of tensile loading (20 cycles at 30% strain amplitude) and subsequent stabilization after 30-50 cycles [67,68], the greater decreases observed in this study are likely associated with the extended number of cycles and the presence of pore structures, which may introduce stress concentrations and facilitate mechanical relaxation. Additionally, no consistent relation was observed between the laser processing parameters and the extent of modulus reduction, suggesting that fatigue-induced softening is governed primarily by the overall membrane architecture rather than fabrication-associated effects. In OoC systems requiring sustained or repeated mechanical stimulation, these findings highlight the importance of designing porous barrier membranes for OoC integration, with consideration for the effects of thin film architecture on membrane’s mechanical behavior.

### 3.3. Membrane surface chemical composition

To assess potential chemical modifications arising from the laser ablation process, Raman spectroscopy was performed on the porous membranes. In particular, attention was paid to the formation of any by-products that may influence cytocompatibility. Two locations within the pore microenvironment were identified for spectral analysis: directly at the pore edge (Figure 3B, Point X) and at a radial distance of 5 μm (Figure 3C, Point Y). Across all fabrication conditions, analysis of the Raman spectra acquired at both measurement locations on the exit surface showed no discernible differences compared to plain PDMS controls (Figure 3A). No additional peaks were detected following laser processing, and the characteristic PDMS molecular vibrational modes (Table 5) were preserved with normalized peak intensities. Similar results were observed for the entry surface (Figure S4), indicating no new chemical entities being created on either side of the membrane.

**Table 5.** The characteristic Raman peaks of PDMS and their corresponding chemical bonds.

| Raman shift ($\text{cm}^{-1}$) | Chemical Bond |
| --- | --- |
| 488 | Si-O-Si symmetric stretching |
| 687 | Si-CH<sub>3</sub> symmetric rocking |
| 709 | Si-CH<sub>3</sub> symmetric stretching |
| 788 | CH<sub>3</sub> asymmetric rocking |
| 859 | CH<sub>3</sub> symmetric rocking |
| 1263 | CH<sub>3</sub> symmetric bending |
| 1411 | CH<sub>3</sub> asymmetric bending |

**Figure 3.**
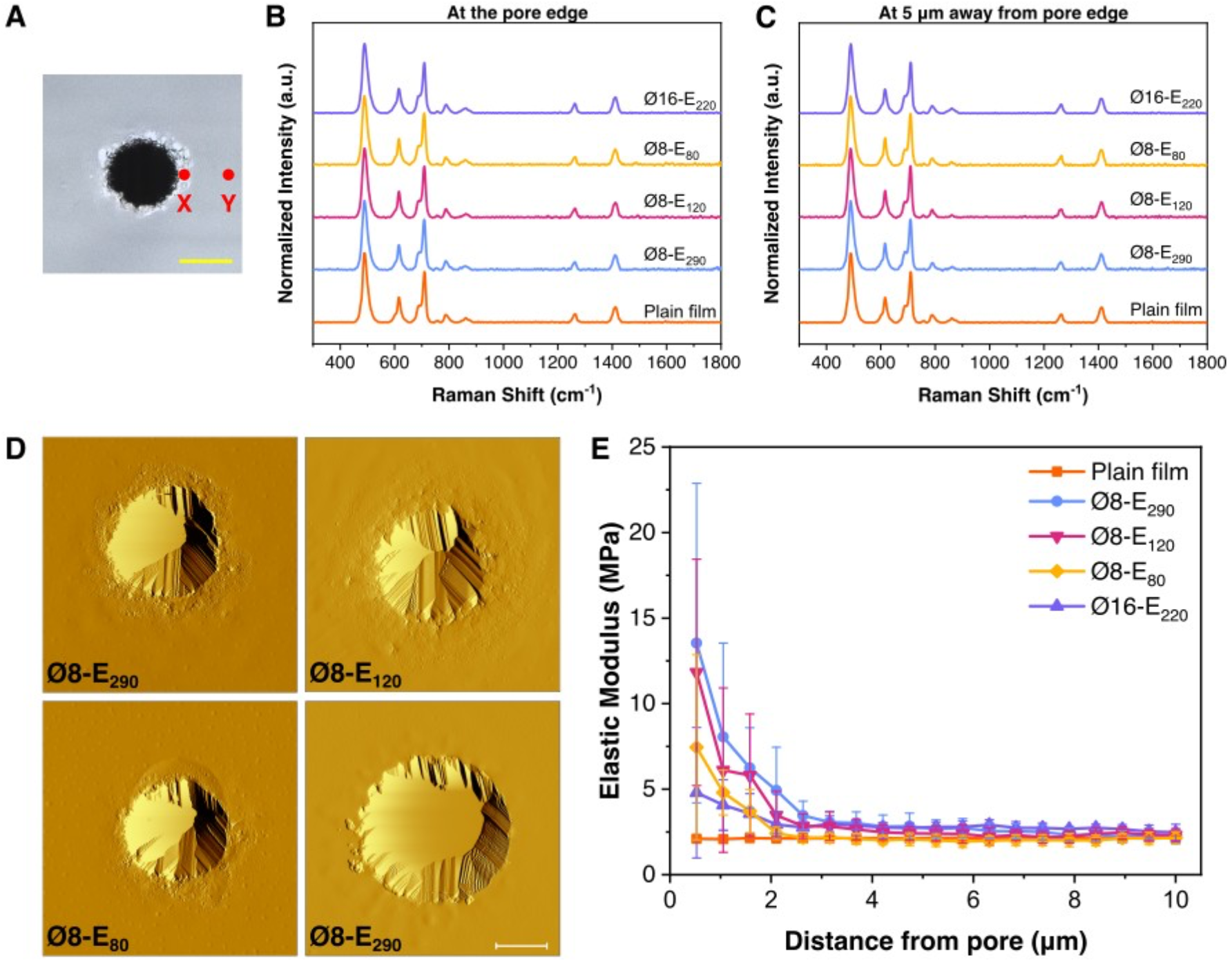
Surface elastic modulus increases near the pore boundary, while chemical analysis confirms no new bonds are generated during laser ablation. (A) Confocal microscopy image of the pore exit surface. The red dots indicate the locations selected for Raman analysis. (scale bar: 5 μm) (B, C) Raman spectra of the PDMS membranes measured (B) at the edge and (C) at 5 μm away from the edge of a pore. (D) AFM topography images of the exit surface of pores fabricated under different laser parameters (scanned left to right). (E) The exit surface Elastic modulus of the PDMS membranes measured around the pore edges. The error bars represent Mean ± 1 SD (n = 3).

Notably, a characteristic crystalline silicon (c-Si) peak at ∼519 cm^−1^, frequently reported in studies involving laser interaction with PDMS, was not observed. This feature has predominantly been associated with nanosecond and visible to near-infrared fs-laser ablation, where photothermal and plasma-mediated effects can induce silicon crystallization [69–71]. Its absence in the present study is consistent with prior reports demonstrating that such signatures are not generated under ultraviolet femtosecond irradiation conditions [71,72]. Therefore, though there might be variations in the PDMS cross-linking density in the immediate areas surrounding the pores, these results suggest that generation of new chemical moieties within the PDMS substrate due to the UV fs-laser ablation process is very unlikely, thus supporting the suitability of the fabricated membranes for cell-interfacing applications.

### 3.4. Surface mechanical properties

Atomic force microscopy (AFM) was employed to characterize the surface mechanical properties of the exit side of the membrane, corresponding to the region intended for cell seeding. Given that the mechanical stiffness of the substrate could influence cellular behavior, local variations in elastic modulus induced by laser processing were evaluated [60].

Analysis of surface topography at the pore exit and surrounding regions (Figure 3D) showed the presence of wrinkles in the immediate vicinity of pore edges, indicating a heat-affected zone generated during laser ablation. However, these features were constrained to the regions close to the pore boundary, with the rest of the surrounding regions appearing relatively smooth. Across all the laser fabrication conditions, an increase in stiffness was observed at the edge of the pore, with modulus values peaking at the pore boundary and progressively declining with increasing radial distance (Figure 3E). This gradient was statistically significant between measurements taken 0.5 µm and 10 µm from the pore edge (p-values included in Table S3), indicating that thermally induced stiffening is concentrated at the ablation boundary rather than being uniformly distributed across the membrane surface. Such localized mechanical modification is consistent with thermally induced post-curing of thermoset PDMS, with the effect decreasing in magnitude as the distance from ablation site increases, reflecting the limited spatial extent of heat transfer in the material.

Among the Ø8 pore membranes, the highest peak modulus was observed for Ø8-E_290_ condition (13.54 MPa), with decreasing modulus recorded for lower energy inputs (11.82 MPa for Ø8-E_120_ and 7.44 MPa for Ø8-E_80_). However, the Ø16-E_220_ condition exhibited a comparatively lower peak elastic modulus of 4.79 MPa despite a higher total pulse energy input. This discrepancy is likely caused by the Gaussian profile of the laser beam’s energy distribution, wherein most of the pulse energy is concentrated within a spot size of 8-10 μm. As a result, compared to larger pores (Ø16), pores with diameters (Ø8) smaller than the laser spot size experience greater thermal energy along the edges, leading to enhanced localized heating and a higher elastic modulus. Around 2-3 μm away from the pore edge, the measured elastic modulus begins to converge to values comparable to plain PDMS (∼2.13 MPa), indicating that the bulk of the exit surface of the membrane remains largely unaffected by the ablation process and mechanically unaltered, suggesting that laser-induced modulus modifications are highly localized.

### 3.5. Cell viability

PDMS was selected as the elastomeric template for membrane fabrication due to its well-established cytocompatibility and its use as a stiffness-tunable cellular substrate for *in vitro* tissue mimicry [11,22,31,32]. To evaluate whether fs-laser fabricated membranes were suitable for cellular interfacing, NIH 3T3 fibroblasts were seeded on fabricated membranes and viability assessed at day 1 and day 3 post-seeding. Fluorescent microscopy confirmed sustained cell viability and active proliferation across all membrane conditions (Figure 4A). By day 3, we observed that the cells adhered to and proliferated along pore perimeters, as the available surface area became limiting, prompting further investigation of the migration potential of cells and small molecules through the porous architecture. Raman spectroscopy conducted at pore edges revealed no new spectral peaks nor shifts in existing peaks relative to spectra acquired 5 µm from pore boundaries on unablated regions (Figure 3B-C) indicating that the bulk chemical composition of the PDMS was preserved during laser processing. The absence of detectable cytotoxicity and the lack of chemically altered surface species together confirm that the fs-laser fabrication does not compromise membrane biocompatibility. These findings are consistent with Berrah, *et al*., who reported that human adipose-derived mesenchymal stem cells remained viable for at least 48 hours post-seeding on fs-laser micro-drilled PDMS membranes, thus corroborating the cytocompatibility of fs-laser ablated membrane fabrication strategy [34]. These results establish that fs-laser drilled porous PDMS membranes are cytocompatible and suitable for applications requiring direct contact with cells and engineered tissue constructs.

**Figure 4.**
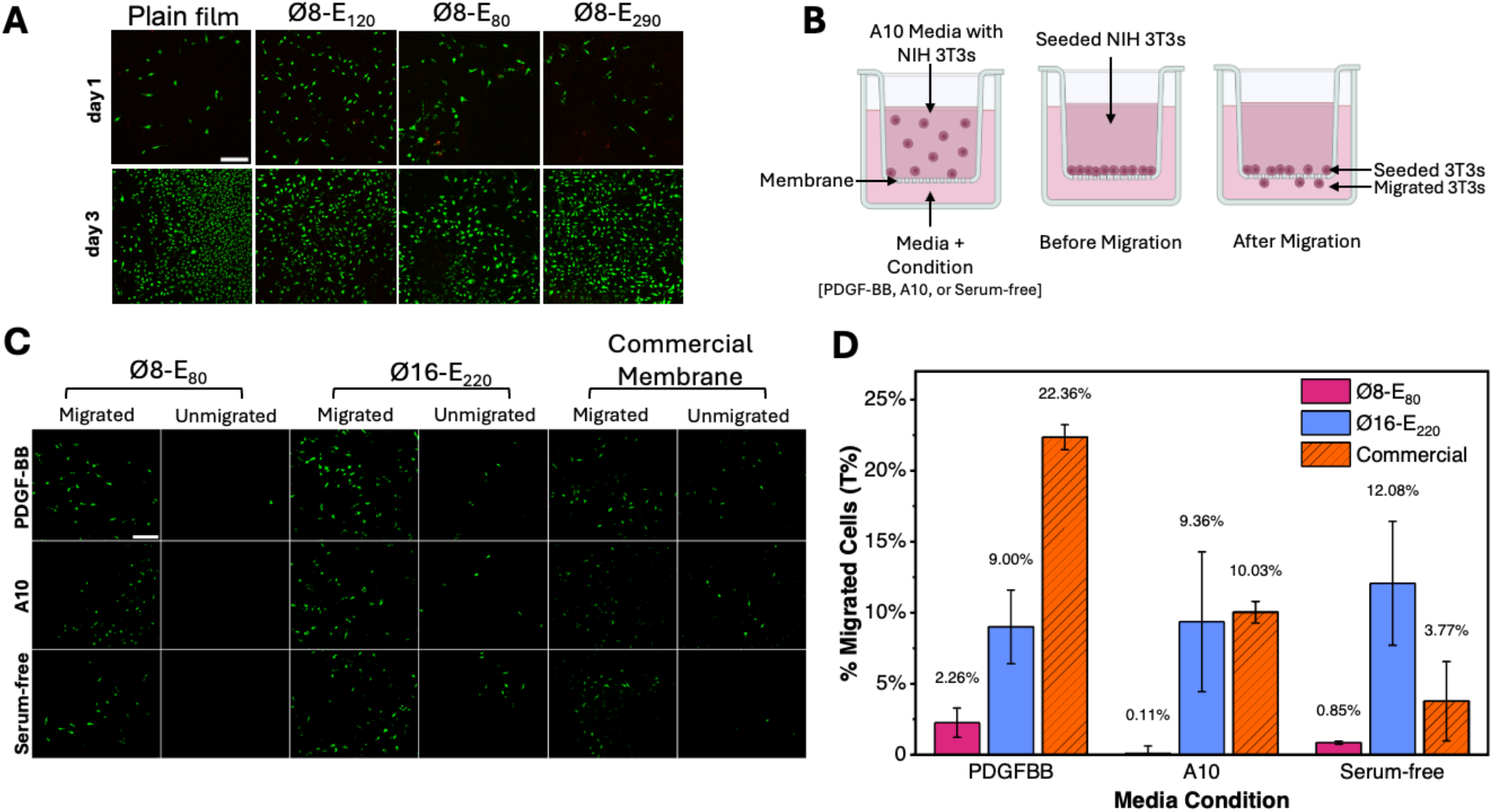
Femtosecond (fs) laser microdrilled membranes maintain viability and limit migration of NIH 3T3s. (A) Representative fluorescent images show live (green, 488nm)/dead (red, 561nm) cells on the control, and on membranes Ø8-E_290_, Ø8-E_120_ and Ø8-E_80_ imaged on day 1 and day 3 (∼80%) in culture. N=3 membranes, n=3 areas/sample. (B) BioRender schematic of the transmigration assay setup illustrating cell seeding, media conditions, and migration across the membrane. (C) Representative images show NIH-3T3 fibroblasts on the unmigrated and migrated surfaces of small-pore (Ø8-E80), large-pore (Ø16-E220), and commercial membranes after 24 h under PDGF-BB, A10, or serum-free (SF) conditions. (D) Percent transmigrated cells (T%) across fabricated PDMS membranes and commercial membrane controls. Data represent mean ± SEM (n = 3 inserts per media condition). Larger pores significantly increased transmigration compared to small pores, and PDGF-BB produced the highest migration across membrane types. Commercial membrane controls exhibited the expected media-dependent trend whereby cell migration in PDGF-BB was the greatest, followed by A10 and the least migration in serum-free medium; fs-laser fabricated PDMS membranes exhibited reduced sensitivity to differences in media composition. All scale bars are equal to 250 μm.

### 3.6. Cell transmigration

Having established membrane cytocompatibility, the functionality of fs-laser-fabricated membranes in regulating cell transmigration was evaluated, as selective barrier function is a critical performance requirement for compartmentalized OoC applications. Transmigration of NIH 3T3 fibroblasts was evaluated across two pore geometries, Ø8-E_80_ (8 µm exit diameter) and Ø16-E_220_ (16 µm exit diameter), in the presence and absence of the chemoattractant platelet-derived growth factor-BB (PDGF-BB). Commercial FluoroBlok™ inserts were used as controls (Figure 4B-D). The two pore diameters were selected to capture the transition regime for confined nuclear migration, as transmigration arrest has been reported when pore cross-sectional area decreases to approximately 10% of the nuclear cross-sectional area [73,74]. Under these conditions, for the NIH 3T3 cells with an average nucleus size of 10-15 µm, when compared to the 16 µm pores, the 8 µm pores present a more restrictive environment.

Transmigration through the laser-fabricated membranes indicated a strong dependence on pore size, with a more variable and condition-dependent response to chemoattractant stimulus. The commercial inserts used for the transmigration assay displayed the expected chemotaxis-dependent hierarchy: transmigration was greatest under PDGF-BB stimulation (22.36%), intermediate in complete A10 medium (10.03%), and lowest in serum-free medium (3.77%). In the laser-fabricated membranes transmigration increased with both larger pore diameter and PDGF-BB stimulation. However, the fabricated membranes diverged from the commercial control in the relative ordering of transmigration observed in A10 versus serum-free medium conditions. Under PDGF-BB, Ø16-E_220_ membranes exhibited greater transmigration than Ø8-E_80_ membranes (9.00% vs. 2.26%). Across the remaining media conditions, pore-size dependency on migration was observable, albeit with greater variability. For Ø8-E_80_ membranes, transmigration remained low in both A10 and serum-free medium (0.11% and 0.85%, respectively), suggesting that geometric constriction might be dominating over soluble cue responsiveness at this pore scale.

For Ø16-E_220_ membranes, serum-free medium produced marginally higher transmigration than A10 medium (12.08% vs. 9.36%). This was in contradiction to the hypothesized trend whereby A10 medium supports greater transmigration than serum-free medium in the laser-fabricated membranes. This deviation may be due to structural differences between the fs-laser fabricated PDMS membranes and the commercial inserts. The PDMS membranes are significantly thicker (50 µm vs. ∼15 µm) with a conical pore architecture, which may introduce additional mechanical and topographical barriers to transmigration, reducing the relative contribution of serum-derived soluble cues. In the thinner commercial inserts, media composition more directly modulates the chemoattractant gradient, yielding the clear media-dependent hierarchy observed in controls. These trends suggest that, within the laser-fabricated system, membrane micro-architecture may play a more influential role than soluble factor availability in governing cellular migration, and the differences in the cellular response across various pore diameters are more likely to be distinguishable in the presence of a chemoattractant in the conditioned media.

Overall, these results demonstrate that fs-laser-fabricated PDMS membranes provide tunable, geometry-dependent regulation of cell transmigration, and that pore size could be leveraged as a practical design parameter for decoupling cell migration from molecular transport in barrier-chip architectures.

### 3.7. Molecular diffusion and selective barrier function

A key design objective of the fabricated membranes was to enable tunable pore geometry to modulate cell migration while still allowing molecular exchange across the compartments. To characterize the molecular transport behavior of the fs-laser fabricated membranes, diffusion of 10 kDa and 250 kDa FITC-dextran as representative proxies for growth factor and glycoprotein molecular weights, respectively, was assessed across Ø8-E_80_ membranes [75,76] (Figure 5A). Time-course fluorescence imaging revealed that both dextran fractions diffused freely across the membranes, reaching equilibrium within 120 seconds of infusion (Figure 5B) [8,77]. This free molecular transport across the membrane was expected, owing to the difference in scale between the pore diameter (6–20 µm) and the dextran tracers (nanometer scale), and is further supported by Lebrun *et al*., who demonstrated uninhibited 10 kDa dextran diffusion even across membranes with pore diameters as small as 200 nm, with diffusivity increasing with the pore size [77,78]. Exponential fitting of diffusion profiles was used to calculate the time (***T***_***c***_) [79] taken for respective dextran sizes to reach 63.2% of their diffusion equilibrium in the system (Figure 5C), yielding time constants of 9.59 s and 53.2 s for 10 kDa and 250 kDa dextran, respectively. The diffusion profiles of the dextran molecules indicate that molecular transport of both the species was not inhibited by the membranes and is likely weight (hydrodynamic diameter)-dependent. The rapid equilibration observed here is important for membrane integration with OoCs, where unimpeded molecular communication across the membranes is necessary to preserve paracrine signaling fidelity and improve the temporal resolution of biomolecule sampling following injurious or inflammatory stimulation [80–84].

**Figure 5.**
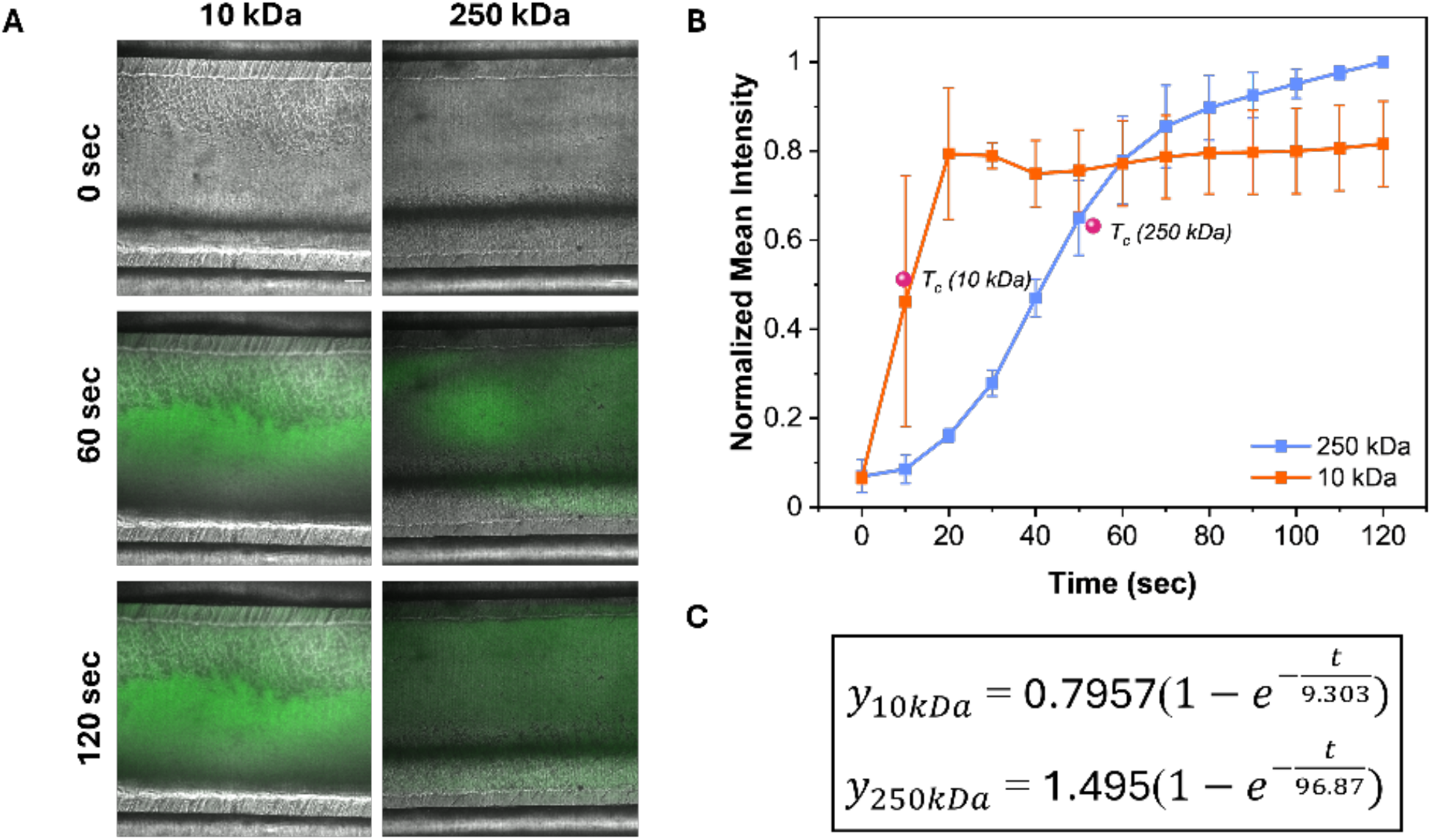
Charged biomolecules readily diffuse through fs-laser patterned membranes. (A) Representative images show fluorescein dextran diffusing through membranes over time (scale bar = 100 µm). (B) Diffusion profile of 10 kDa (orange) and 250 kDa (blue) fluorescein dextran plotted as normalized intensity over time for fs laser micropatterned membranes. (C) Diffusion time constants represent the time it takes for respective dextran sizes to reach 63% of their equilibrium. n=3 experiments.

Rapid molecular diffusion alone, however, is insufficient to achieve physiologically relevant compartmentalization in OoC systems. Many OoC architectures rely on porous membranes to maintain physical separation of distinct cell populations without eliminating biochemical communication and compromising barrier function. This functional duality of molecular permeability alongside cellular compartmentalization is central to barrier-chip architectures, such as lung alveolar-capillary and gut barrier models, in which porous membranes sustain discrete microenvironments while permitting biochemical exchange [4,82]. The current results demonstrate that fs-laser-fabricated PDMS membranes satisfy both requirements simultaneously. While 10 kDa and 250 kDa dextran diffused rapidly and without restriction across Ø8-E_80_ membranes, fibroblast transmigration through the same geometry was constrained, even under PDGF-BB stimulation. Increasing pore diameter to 16 µm enabled increased cell migration, thus suggesting that pore geometry could be an effective lever to modulate cellular versus molecular permeability within the same membrane.

This decoupling of molecular transport from cell migration has direct implications for OoC design. Applications requiring long-term compartment integrity, such as epithelial or endothelial barrier models, are well served by the cell-restrictive behavior of the smaller pore geometry, while models incorporating active transmigration events, such as immune cell extravasation or recruitment assays, can be accommodated by transitioning to larger pore geometries [85]. Together, these findings establish the fs-laser-fabricated PDMS membranes as tunable selective interfaces capable of maintaining rapid molecular communication while preserving cellular compartmentalization and providing controlled cellular exchange, making them a versatile component of OoC systems.

## 4. Conclusion

This study presents a two-stage fs-laser fabrication process for producing drilled microporous PDMS membranes for OoC applications. The relationship between process parameters, membrane geometry, mechanical properties, surface chemistry, and biological performance was evaluated. The findings demonstrate that the fabrication approach affords control over pore geometry while preserving the chemical integrity and cytocompatibility of the substrate, establishing a route toward soft, tunable, porous membrane interfaces for compartmentalized micro-physiological systems.

Pulse repetition frequency (PRF) was observed to be a dominant parameter governing entry diameter and surface roughness across both fabrication stages, while exit diameter was primarily determined by PRF and NP at the first stage. The development of a predictive model also demonstrated the capability of the two-stage process to modulate the entry and exit-end diameters independently. In the long term, this enables the tunable design of conical pore architecture without iterative experimental optimization. The elastic moduli of the laser patterned membranes, observed across all fabrication parameters, was 25% higher than that of plain, non-patterned PDMS. The mechanical stiffness, as examined by AFM and nanoindentation, increased within micron-scale vicinity of pore boundaries and decreased with increasing radial distance, exhibiting a gradient that may influence cellular behavior at the pore interface.

Fatigue testing revealed a 5 to 12% reduction in elastic modulus following 500 cycles of 30% strain, providing initial data to inform device lifetime estimates for OoCs integrated with fs-laser fabricated porous membranes, particularly under physiologically and pathologically relevant loading conditions. Raman analysis of the regions surrounding the pores showed that characteristic PDMS peaks retained their normalized intensities and no new peaks emerged, indicating that laser ablation did not introduce new chemical bonds and that the membrane surface composition was largely unaltered, consistent with the cytocompatibility outcomes observed with NIH 3T3 cells.

Pore size was the primary determinant of fibroblast transmigration, with smaller geometries functioning as effective cellular barriers even under chemoattractant stimulation, while molecular transport remained rapid and uninhibited across the same geometries. The decoupling of cellular and molecular permeability through the appropriate selection of pore geometry emerged as a key functional attribute of the fs-laser patterned membranes, as it provides an independent and scalable design parameter for constructing multi-tissue compartmentalized systems. These properties are particularly relevant to joint-on-chip applications, where existing models achieve either biochemical crosstalk between synovial compartments [86,87] or mechanically driven osteoarthritic phenotypes [88], but none integrate mechanical stimulation with multi-tissue biochemical exchange simultaneously. The membranes developed in this study offer a practical route towards bridging this gap, enabling soluble mediators from mechanically loaded tissue compartments to diffuse across selective interfaces while cell passage remains controlled.

In conclusion, the two-stage fs-laser fabrication approach enables the design and development of laser-drilled, microporous PDMS membranes with tunable pore geometry, resilient mechanical behavior, and cytocompatibility. Furthermore, the membranes allow for rapid biomolecular cross-talk while preserving cellular compartmentalization via confined pore geometries. These properties establish fs-laser fabricated membranes as selective and adaptable interfaces for compartmentalized OoC systems, particularly relevant for joint-on-chip architectures where integrated mechanical stimulation alongside multi-tissue, multi-compartment biochemical exchange remains an unmet design challenge.

## Acknowledgements

This work was funded by National Institutes of Health (NIH) grant R01 AR083379.

## Author Contributions

All authors contributed to the conceptualisation and design of the study. AG, GW developed the methodology for laser-related fabrication and physico-chemical characterization; AG, AJD carried out the experimental investigation and performed data analysis. SG, IR, SES, CN developed the methodology for biological charcaterization and molecule transportation assays; SG, IR carried out the corresponding experimental investigation and performed data analysis. AH, HK, ZC assisted in the operational optimization of laser patterning set-up. AG, AJD, SG, IR wrote the original draft of the manuscript. AJD, SES, CN, GW reviewed and edited the manuscript. AG, AJD, SG, IR were responsible for visualisation. KJ supported project execution by providing the laser set-up used in the study. GW, CN supervised the study, and were responsible for project administration and funding acquisition. All authors have read and agreed to the published version of the manuscript.

## Conflicts of Interest

The authors declare no conflict of interest.

## Notes

### Competing Interest Statement

The authors have declared no competing interest.

