## Supplementary Information for "Multi-step femtosecond laser-fabricated membranes for regulated migration of biomolecules and cells"

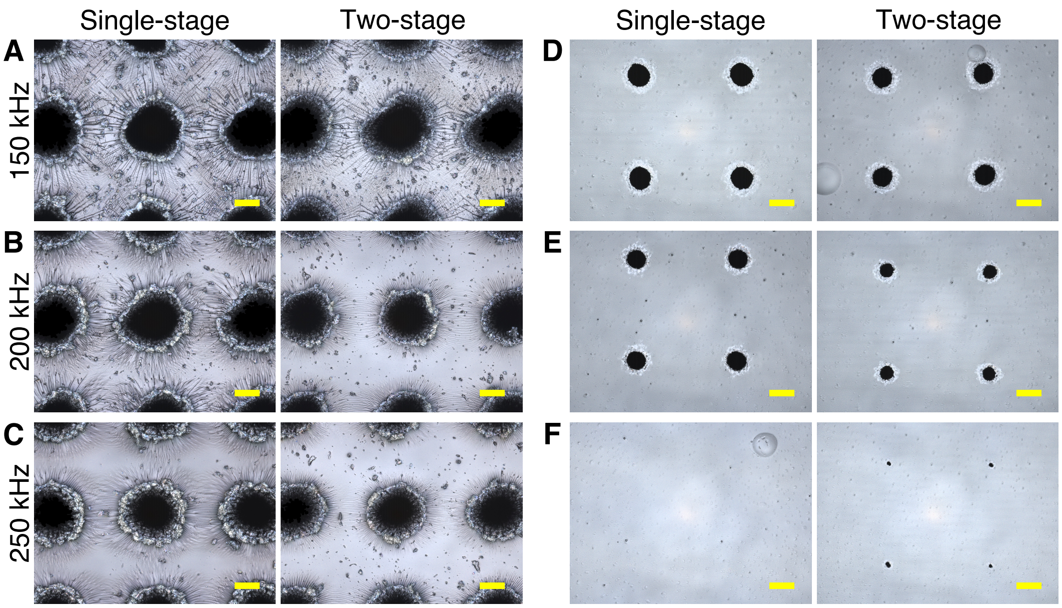


**Figure S1.** Thermal damage on the entry surface is lower in membranes fabricated using a two-stage fs-laser ablation process. (A-C) Confocal microscopy images showing entry surface morphologies of the single-stage (40 pulses) and two-stage (20+20 pulses) fabricated membranes at 150, 200 & 250 kHz laser PRFs. (D-F) Corresponding exit surface morphologies.

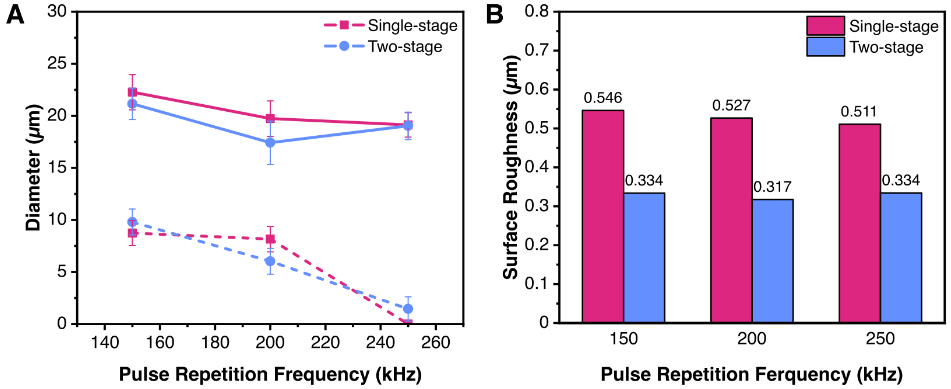


**Figure S2.** Two-stage laser fabrication reduces entry surface roughness without heavily altering pore diameters. (A) Evolution of pore entry (solid lines) and exit (dotted lines) diameters with laser PRF in single- (40 pulses) and two-stage (20+20 pulses) fabrication processes. (B) Corresponding entry surface roughness of membranes fabricated using single- and two-stage processes.

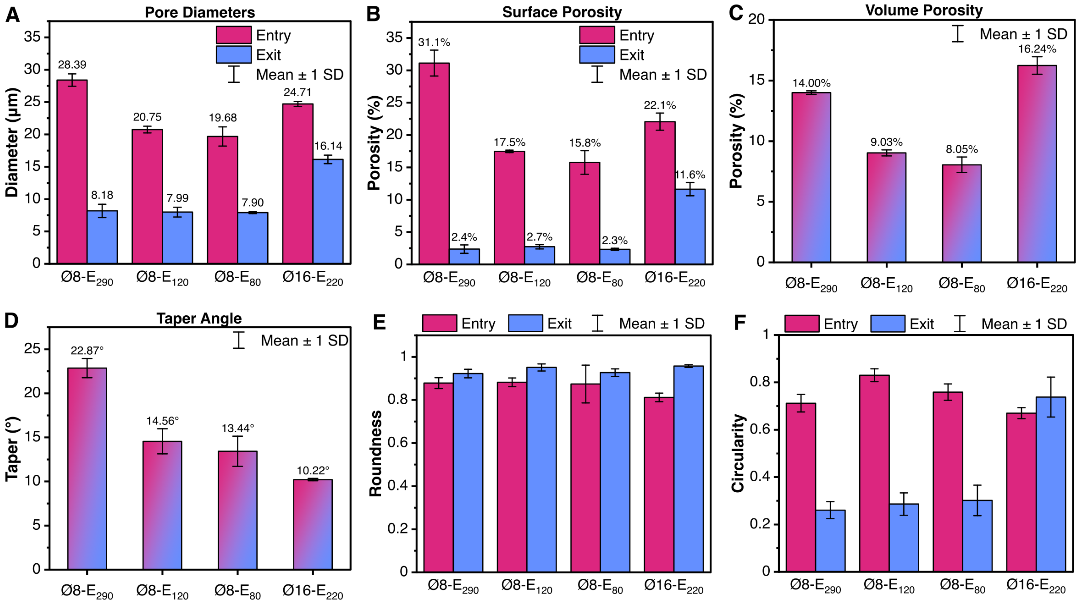


**Figure S3.** Pore dimensions for the fs-laser fabricated membranes

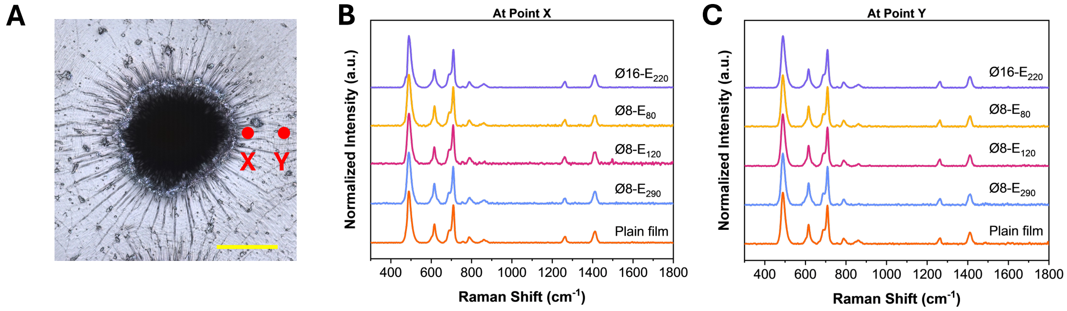


**Figure S4.** Raman analysis on the entry surface of fs-laser fabricated membranes. (A) Confocal microscopy image of the pore entry surface. The red dots indicate the locations selected for Raman analysis. (scale bar: 5 µm) (B, C) Raman spectra of the PDMS membranes measured (B) at the edge and (C) at 5 µm away from the edge of a pore.

**Table S1.** Results of experiments with laser parameters and membrane characteristics.

| I - PRF | I - NP | II - PRF | II - NP | Exit Dia (μm) | Entry Dia (μm) | Entry $S_q$ (μm) |
|---|---|---|---|---|---|---|
| 100 | 25 | 100 | 40 | 15.597 | 32.703 | 1.269 |
| 300 | 25 | 150 | 20 | 0 | 20.683 | 0.34 |
| 100 | 40 | 300 | 20 | 9.767 | 33.094 | 0.751 |
| 100 | 20 | 250 | 20 | 0 | 28.078 | 0.454 |
| 100 | 40 | 100 | 40 | 20.689 | 37.254 | 0.894 |
| 250 | 20 | 100 | 25 | 0 | 29.822 | 0.725 |
| 300 | 40 | 100 | 20 | 0 | 29.72 | 0.582 |
| 150 | 25 | 300 | 25 | 5.566 | 23.866 | 0.393 |
| 150 | 20 | 150 | 35 | 14.234 | 25.158 | 0.612 |
| 100 | 20 | 100 | 20 | 0 | 32.289 | 0.841 |
| 150 | 40 | 250 | 40 | 16.186 | 24.834 | 0.726 |
| 300 | 20 | 300 | 40 | 0 | 21.915 | 0.256 |
| 150 | 35 | 100 | 20 | 13.687 | 31.919 | 0.529 |
| 250 | 40 | 300 | 20 | 4.198 | 21.664 | 0.33 |
| 100 | 40 | 150 | 25 | 15.722 | 35.822 | 1.01 |
| 300 | 40 | 300 | 40 | 4.036 | 25.183 | 0.349 |
| 250 | 25 | 250 | 35 | 6.624 | 23.166 | 0.177 |
| 250 | 35 | 150 | 40 | 14.39 | 26.64 | 0.268 |
| 300 | 20 | 300 | 20 | 0 | 17.735 | 0.148 |
| 300 | 35 | 250 | 25 | 3.549 | 20.906 | 0.21 |
| 100 | 20 | 300 | 40 | 3.418 | 29.38 | 0.521 |
| 100 | 35 | 300 | 35 | 9.97 | 32.852 | 0.643 |
| 300 | 40 | 100 | 35 | 6.758 | 31.968 | 0.547 |
| 300 | 20 | 100 | 40 | 0 | 31.667 | 0.648 |
| 300 | 20 | 100 | 20 | 0 | 28.285 | 0.492 |

**Table S2.** ANOVA results with contribution ratios for exit Diameter, entry Diameter and entry surface roughness.

| **Exit Diameter** | | | | | |
|---|---|---|---|---|---|
| Factor | DF | Sum of Squares | F-Ratio | p-value | Contribution (%) |
| I-PRF | 3 | 331.702 | 8.563 | 0.0026 | 41.03 |
| I-No. of Pulses | 3 | 243.512 | 6.287 | 0.0083 | 30.11 |
| II-PRF | 3 | 89.590 | 2.313 | 0.1278 | 11.08 |
| II-No. of Pulses | 3 | 143.72 | 3.71 | 0.0425 | 17.76 |
| **Entry Diameter** | | | | | |
| Factor | DF | Sum of Squares | F-Ratio | p-value | Contribution (%) |
| I-PRF | 3 | 266.633 | 18.34 | <0.0001 | 45.25 |
| I-No. of Pulses | 3 | 54.351 | 3.739 | 0.0417 | 9.22 |
| II-PRF | 3 | 242.964 | 16.72 | 0.0001 | 41.24 |
| II-No. of Pulses | 3 | 25.238 | 1.736 | 0.2127 | 4.28 |
| **Entry Surface Roughness** | | | | | |
| Factor | DF | Sum of Squares | F-Ratio | p-value | Contribution (%) |
| I-PRF | 3 | 0.813 | 13.429 | 0.0004 | 51.53 |
| I-No. of Pulses | 3 | 0.122 | 2.009 | 0.1665 | 7.71 |
| II-PRF | 3 | 0.560 | 9.248 | 0.0019 | 35.49 |
| II-No. of Pulses | 3 | 0.083 | 1.374 | 0.298 | 5.27 |

**Table S3.** Results of the Welch's independent samples t-test performed between elastic modulus measurements at the pore edge (~0.5 μm) and at ~10 μm away from pore edge, showing significant difference.

| Membrane | n | Elastic Modulus (MPa) (M ± SD) | | DF | t Ratio | p | d |
|---|---|---|---|---|---|---|---|
| | | At 0.5 μm | At 10 μm | | | | |
| Ø8-$E_{290}$ | 25 | 13.54 ± 9.73 | 2.25 ± 0.42 | 24.09 | −5.80 | <.0001 | 1.64 |
| Ø8-$E_{120}$ | 15 | 11.83 ± 7.07 | 2.19 ± 0.39 | 14.09 | −5.20 | <.0001 | 1.9 |
| Ø8-$E_{80}$ | 15 | 7.45 ± 5.78 | 2.15 ± 0.38 | 14.12 | −3.54 | .003 | 1.29 |
| Ø16-$E_{220}$ | 25 | 5.33 ± 3.32 | 2.51 ± 0.64 | 25.8 | -4.17 | 0.0003 | 1.18 |
